# Recognition in Confinement: The Dynamics of Homologous Gene Pairing

**DOI:** 10.64898/2026.01.10.698804

**Authors:** Ehud Haimov, Jonathan G. Hedley, Natalia E. Simonowicz, Yaxuan Xiao, Andrew Stannard, Gleb Oshanin, Angelo Rosa, Yuval Elani, Alexei A. Kornyshev

## Abstract

How can protein-independent, side-by-side alignment of homologous double-stranded DNA (dsDNA) arise as an early step in genome repair and genetic exchange, and what determines their kinetics under confinement? Although recombination proteins are known to facilitate this process, experiments indicate that an initial side-by-side arrangement of homologs can occur even in their absence. Helical Coherence Theory (HCT) proposes that sequence-dependent distortions of the double helix lead to commensurate charge patterns between interacting dsDNAs, favouring homologous over non-homologous alignment. However, HCT has largely been developed for straight, rigid dsDNA rods, an approximation limited to roughly one persistence length (*ℓ*_*B*_ ≈ 50 nm), leaving the question open as to how recognition manifests in longer, fluctuating chains. Moreover, the dynamics and timescales of homolog searching under confinement, and the internal dynamics of the paired state, such as transient formation of local unpaired regions (“bubbles”) and end fraying, remain unexplored. Here we extend HCT to dsDNA chains of multiple persistence lengths under confinement using coarse-grained simulations with a custom HCT-based force field, in order to come closer to understanding how homologous genes pair *in vivo*. We simulate two homologous chains within spherical cavities of radii 0.3–1 times their free-space gyration radius under two limiting ionic conditions. We found that confinement warrants homologous pairing; it occurs on microsecond timescales and depends non-monotonically on confinement size, with an optimal confinement that accelerates pairing while avoiding tangled states and metastable trapping in an ion-dependent manner. To resolve pairing dynamics, we develop a kinetic theory and fit it to simulation trajectories, distinguishing rates of independent pairing modes underlying bubble formation and fraying. Together, these results support a physical mechanism for homologous pairing beyond the rigid-rod limit and suggest how confinement can promote both encounter and stabilisation within the HCT framework.

## I. INTRODUCTION

The exchange of homologous genes in genetic recombination is a key process in meiosis and DNA repair. It speeds up evolution, enhances genetic diversity and underpins the transfer of genetic material between different bacteria and virus species through horizontal gene transfer [1, 2]. While the protein-mediated stages of recombination are well understood [3], the initial step, i.e. how homologous sequences identify one another to enable precise recombination at corresponding loci, remains unresolved, even in light of recent reports proposing a potential role for cohesin in the early stages of the recombination process during DNA repair [4, 5]. Intriguingly, experimental evidence shows that homolog recognition between genomic regions with high sequence similarity can occur even in the absence of proteins *in vitro*, in simple physiological solutions [6–11].

While the origin of early homology recognition remains unresolved [12–14], several theoretical models support the possibility of protein-independent sequence recognition [15– Recent experimental work [11] has substantiated one such mechanism based purely on the innate physical properties of DNA [18], supporting the view that homology recognition is an inherent, and potentially universal, feature of double helical DNA [25]. This physical recognition mechanism, dubbed Helical Coherence Theory (HCT) rests on the well-established fact that DNA is not an ideal double-helix; rather, its local structure exhibits sequence-dependent distortions of structural characteristics such as helical twist and rise [26– When two rigid double-stranded DNA molecules of identical or nearly identical (homologous) sequences are brought into a 1-to-1 parallel alignment, they can rotate about their long axes to adopt a mutual azimuthal orientation that aligns the negatively charged phosphate backbones of one molecule with the positively charged lines of adsorbed counterions in the grooves of the other (see Fig. 1a). In particular, homologous sequences are considered to be perfectly structurally correlated with one another, and so can maintain this attractive register throughout their length of juxtaposition, thus forming an ‘electrostatic zipper’ [32] between the DNAs. When the degree of phosphate-charge compensation by adsorbed counterions, *θ*, is sufficiently large (typically above 70-75%, i.e. *θ* ≳ 0.75), this effect produces net attraction between the molecules at short interaxial distances; at lower charge compensations, it reduces repulsion. Formally, the electrostatic interaction energy per unit length between two such helices at separation *R* and relative azimuthal angle *δϕ* is

**FIG. 1.**
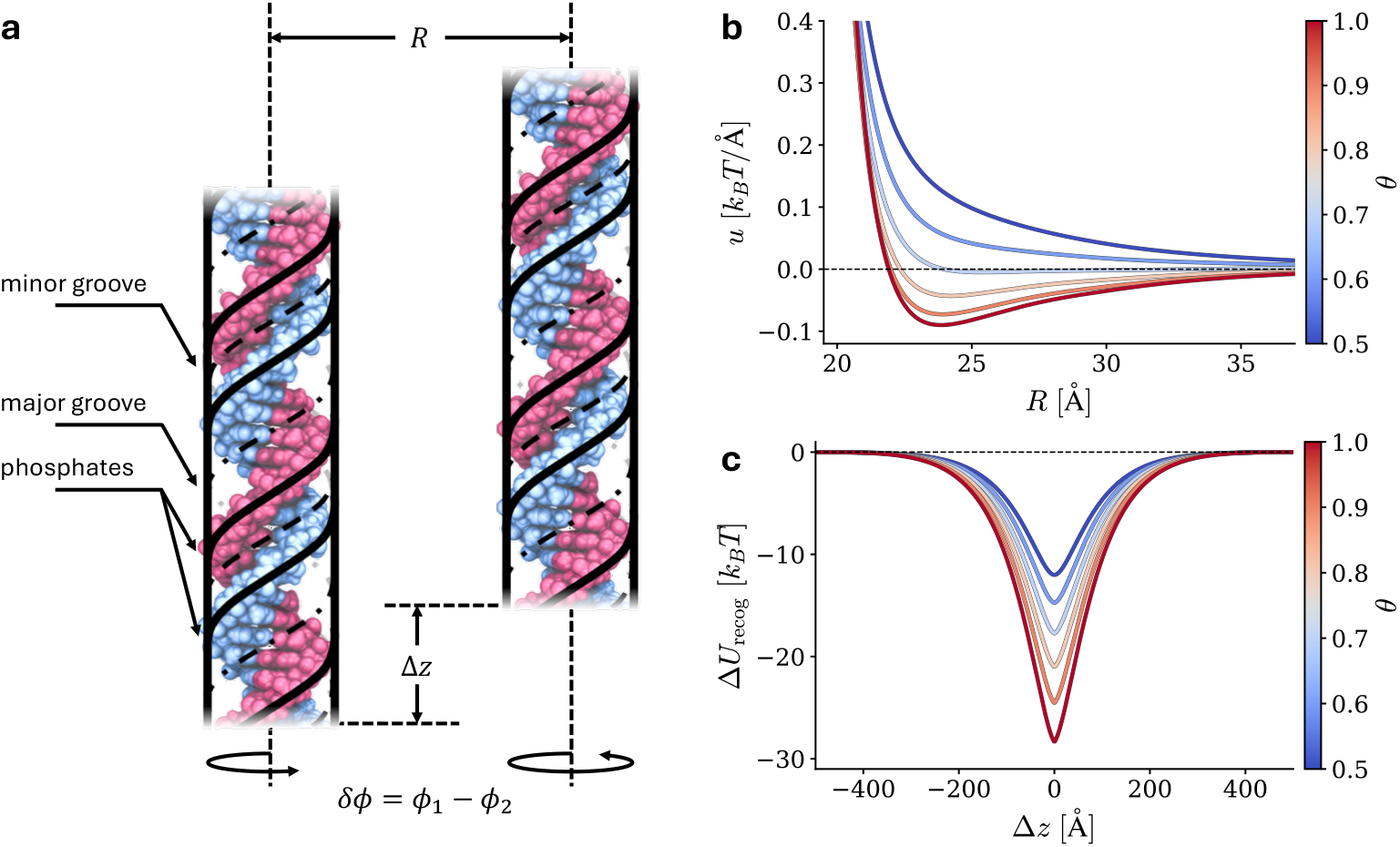
Helical Coherence Theory. (a) Schematic of two parallel DNA duplexes at an interaxial separation *R*, relative azimuthal orientation *δϕ* = *ϕ*_1_ − *ϕ*_2_, and axial shift Δ*z*; the helical charge patterns in the theory arise from the negatively charged double-helical phosphate backbone, and the positively charged lines of condensed counterions in the minor and major grooves. (b) Electrostatic interaction energy per unit length, *u* (Eq. 1), minimised w.r.t. *δϕ*. Negative energy indicates attraction; curves illustrate the progressive transition from repulsion to attraction with increasing charge compensation *θ*. (c) Recognition energy well, Δ*U*_recog_ (Eq. 2) for a juxtaposition window of length *L* = 1000 Å, at a fixed interaxial separation of *R* = 30 Å, minimised w.r.t. *δϕ*. Here, *f*_1_ = 0.2, *f*_2_ = 0.8, *λ*_*c*_ = 100 Å, the Debye length *κ*^−1^ = 7 A, the DNA radius *a* = 10 Å, the helical pitch of DNA, *H* = 34 Åand the dielectric constants of water and the DNA core are *ε* = 80 and *ε*_*c*_ = 2 respectively.

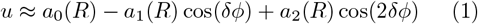

where expressions for the helical harmonics, *a*_*n*_(*R*), are given in the Appendix. Here, *a*_0_ describes a purely repulsive term with two contributions: (i) long range monopole interaction, and, (ii) interaction of helical charges with image charges due to difference in the dielectric constant inside and outside the dsDNA hydrophobic core. *a*_1_ and *a*_2_ are higher harmonics which account for the double helical charge patterns pertaining to the DNA structure. The relative azimuthal orientation that minimises the energy (the ‘zipper’ orientation), is then given by *δϕ*_∗_ = ± arccos(*a*_1_*/*4*a*_2_). Plotting the minimal interaction energy as a function of axial separation *R* in Fig. 1b, it is clear that *θ* modulates the behaviour of the two molecules between repulsion and attraction via the relative contributions of the helical harmonics, with a stable interaxial distance (global minimum energy) forming for *θ* ≳ 0.7.

When two rigid and parallel homologous dsDNAs are shifted along their main axis, by some Δ*z* relative to their favourable 1-to-1 configuration, they gradually lose correlation in sequence-dependent variations in base-pair twist and rise. The loss of registry with Δ*z*, produces a “homology-recognition potential well” in the interaction energy, with half-width *λ*_*c*_ [25], the characteristic shift length over which correlations are lost, called the *helical coherence length* [33]. This is a universal, mesoscopic, characteristic of DNA, being first predicted theoretically [18] and later extracted from classical X-ray fibre diffraction experiments [34], found to be ~ 11 nm [35]. In the case of shifted homologs, Eq. 1 no longer describes the correct DNA-DNA interaction, and the correct formula is given by [25]:

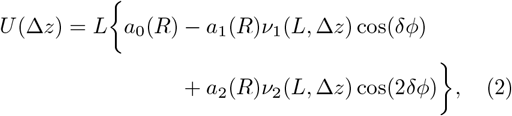

where the recognition coefficients are

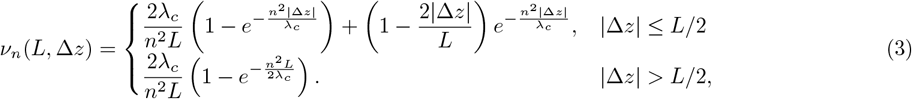

where *L* is the interaction length along which the two DNA molecules are juxtaposed in parallel and at close proximity interaxial distance *R*. The relative azimuthal angle *δϕ*_∗_ that minimises the energy is now given by *δϕ* = ± arccos(*a*_1_*ν*_1_*/*4*a*_2_*ν*_2_). The interaction energy between shifted homologs has a funnel-shape profile, characterised by half-width length of *λ*_*c*_ and a depth set by the sequence-correlation strength and ionic conditions, thus quantifying the crossover from homologous registry at Δ*z* = 0 to effective heterology for Δ*z* ≫ *λ*_*c*_ (see Fig. 1c for the profile of energy difference between those two cases). At a fixed *R*, increasing *θ* amplifies *a*_1_ and *a*_2_, thus deepening this funnelled-shape recognition well.

Several studies suggest that crowding agents can promote homologous pairing. In particular, Baldwin et al. [7] studied how homologous dsDNA oligomers pair under low charge neutralisation, with gradually increasing osmotic pressure via added PEG. For mixtures of two distinct dsDNA sequence populations, domain segregation was observed despite net electrostatic repulsion between duplexes. Complementary evidence comes from single-molecule magnetic-tweezers experiments, which showed that homologous pairing can occur in simple electrolytes, but is accelerated in the presence of crowding agents [8]. From a theoretical perspective, it has also been shown that homologous sequences aligned in an anti-parallel fashion interact no differently from heterologous sequences; in liquid crystalline assemblies, osmotic stress can drive a transition from anti-parallel to parallel alignment, thereby restoring recognition [36]. Together, these observations indicate that crowding-induced compression, and the associated confinement of available configurations, can play a central role in homologous pairing. In cellular environments, analogous compressive effects arise naturally from macromolecular crowding and spatial confinement.

Long homologous dsDNAs, spanning many persistence lengths, have been shown experimentally to form paired structures under physiological salt conditions [8, 9]. These observations were further substantiated by a recent theoretical study based on HCT, which predicted the spontaneous formation of partially paired homologous structures, with localised “bubbles” of unpaired dsDNA between paired regions [37].

Despite the *in vitro* evidence of homologous gene pairing, the mechanism of this process *in vivo* is still debated. One potentially significant factor, that has not yet been explored, is confinement, which in cells can be mediated by membranes or biomolecular condensates. It induces a compressive effect on long dsDNA, altering its motion and interactions. To elucidate the mechanism of homologous pairing in living system, here, we systematically investigate for the first time, two central aspects of homologous pairing of long dsDNA under confinement:

i. the equilibrium and quasi-equilibrium statistics of pairing, in which homologous molecules adopt paired configurations that are stable over long timescales;
ii. the dynamics of the pairing process.

By combining coarse-grained computer simulations based on homology-correlation theory (HCT) with kinetic modelling, we demonstrate that confinement has a dramatic impact on homologous pairing and identify previously overlooked factors that may critically influence the pairing behaviour of homologous genes.

## II. MODELLING APPROACH

### A. Coarse-Grained Description

To simulate two homologous DNA chains confined within a sphere, each double-stranded DNA was coarsegrain modelled as a chain of spherical beads of diameter ℓ_0_ = 2 nm, the value of which is chosen to reflect the realistic width of the molecule. Our coarse-grained model is described in Fig. 2. To capture the mechanical extensibility and large-scale conformational properties of the polymer, we implemented a set of standard mechanical interactions between neighbouring beads, calibrated to reproduce the polymer’s elastic response and gyration radius (see Appendix A.2. for details).

**FIG. 2.**
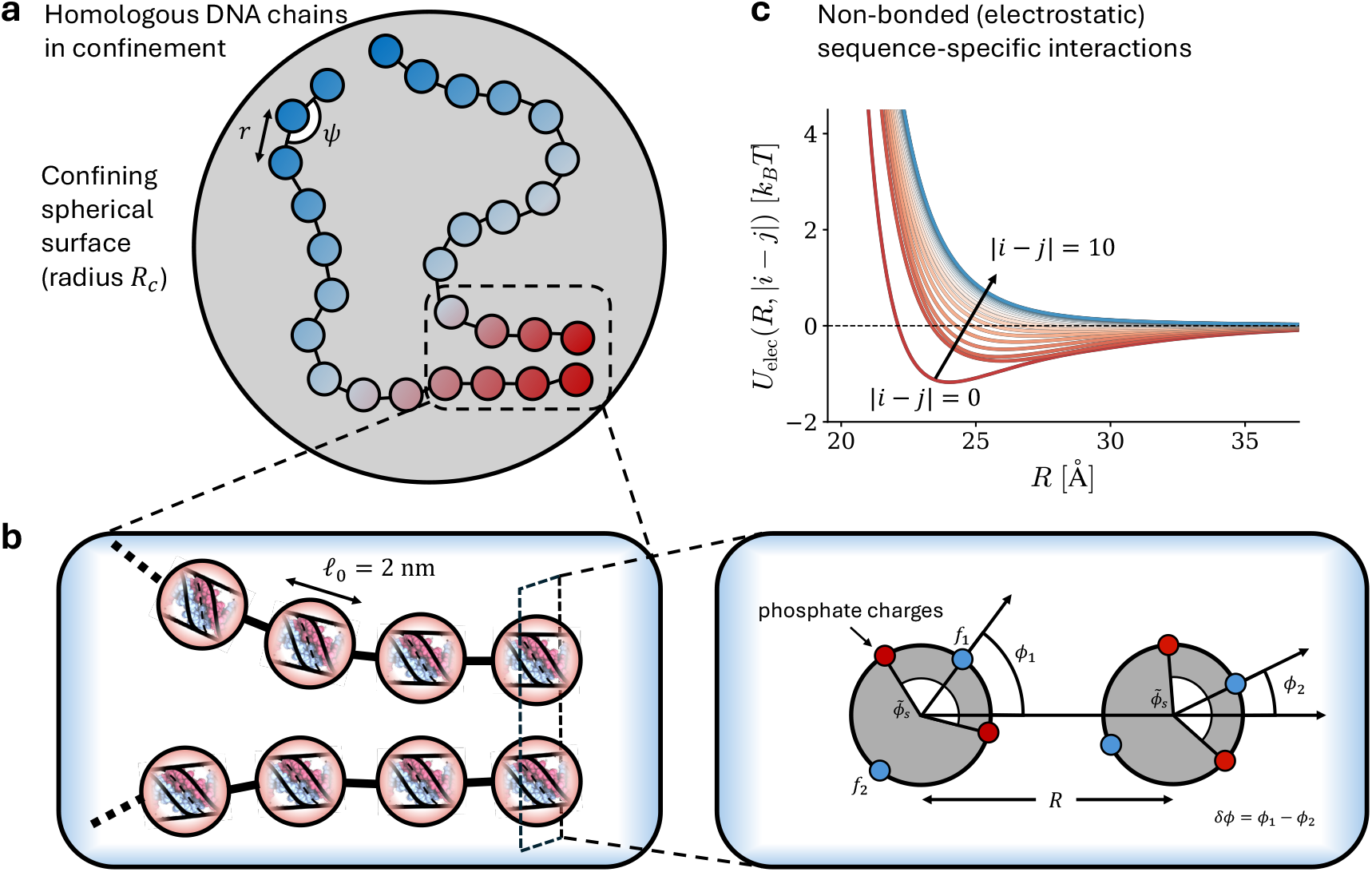
Coarse-grained model of confined DNA. (a) Two homologous DNA molecules are enclosed inside a spherical confinement of radius *R*_*c*_. Each molecule is represented as a series of beads with equilibrium bond length *r* and bond angle *ψ*; bead colours grade from blue to red to denote sequence position, so matching colours identify homologous base-pair registers. (b) Enlarged view of the coarse-grained representation. Each bead corresponds to ℓ_0_ = 2 nm (~ 6 bp) of double-stranded DNA. Non-bonded interactions are computed from an analytical electrostatic model in which helical lines of phosphate charges (red) and partially neutralising counterions (blue) populate the major (*f*_2_) and minor (*f*_1_) grooves. The pair potential depends on the centre-centre distance *R* and the azimuthal offset *δϕ*; the latter is minimised for every pair to obtain the effective interaction. (c) Electrostatic pair potential *U*_elec_(*R*, | *I* − *j*|) for selected sequence offsets |*i* − *j* |, computed from (Eq. 4). Perfectly aligned homologous segments (|*i* − *j*| = 0, red) experience a short range attraction, whereas non-homologous pairs ( |*i* − *j* |≳ 5, blue) are purely repulsive. Intermediate offsets yield progressively weaker attraction, illustrating how sequence specificity is encoded in the interaction.

To keep track of similar (homologous) sequences along the chain and to model their interactions accordingly, we numbered the beads of each DNA molecule. For two overall homologous chains, the electrostatic interaction between bead “*i*” in the first chain and bead “*j*” in the second chain was modelled to reproduce the results of HCT [25] (Eq. 2), in the limiting case of large interaction length *L* ≫ *λ*_*c*_ (see Appendix A.3 for justification):

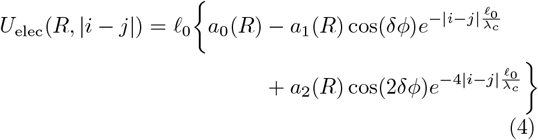

where *R* is the centre-to-centre separation between beads, *δϕ* ≡ ϕ_i_ − *ϕ*_*j*_ where *ϕ*_*i*_ and *ϕ*_*j*_ are the angles stretched between the interaxial line of the sections represented by beads *i* and *j* and the middle of their minor grooves. The functions *a*_0,1,2_(*R*) are the first few dominant harmonics of the electrostatic interaction between two parallel homologous DNA in register, depending on the helical charge patterns formed by the phosphate strands and condensed counterions. These manifest in the set of parameters, 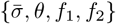, where 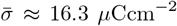 is the average DNA charge density, *θ* is the fraction of DNA charge compensated by condensed counterions, and *f*_1_, *f*_2_ are the fractions of these counterions condensed in the minor and the major grooves respectively (see Appendix A.3. for explicit expressions). The first and second harmonic coefficients are also modulated by:

1. cosine functions of a multiple of the difference in the local angular azimuthal orientations of the DNA segments represented by the beads and,
2. exponentially decaying functions, which account for the effects of an interaxial shift from a 1-to-1 register of the two homologous sections.

The modelling of homologous interactions as presented in Eq. 4 was enabled by the fact that the electrostatic helical coefficients *a*_0,1,2_ decay at least as fast as Debye length ( ~ 7 Å in cytoplasm) – actually faster, with a decay length related also to the helical pitch [19, 20, 32], whereas the homology prefactors decay exponentially with a characteristic length scale which is at least not smaller than *λ*_*c*_*/*4 ≈ 2.5 nm. The two length scales are separated by almost an order of magnitude and can be thus considered independently. In Eq. 4, for each pair of beads, we determined *δϕ* by adopting an adiabatic approximation in which torsional degrees of freedom about the DNA long-axis relax instantaneously to their minimum energy configuration. Therefore, *δϕ* is also a function of distance *R* between the beads (see Appendix A.3).

The DNA molecules were encapsulated within a confining spherical surface of radius *R*_*c*_. The interaction between a bead and the confining wall at a surface-to-surface distance *x* was taken as steric or hydration type force of the form:

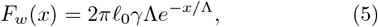

where *γ* ≈ 10^8^ Jm^−3^ [38] is the characteristic repulsion energy per unit volume and Λ ≈ 0.3 nm is the diameter of a water molecule. The relevant geometries for this interaction were accounted for in Eq. 5 by applying a Derjaguin approximation [38].

The displacement of each bead, over each time step, was governed by Langevin dynamics which includes a viscous drag force and a thermal-noise random force, as well as non-random forces (mechanical and electrostatic). The viscous drag force was evaluated using Stokes’ law corresponding to the bead’s size and the viscosity of the solvent. The self-correlation magnitude of the random force was evaluated in a standard way using the fluctuation–dissipation theorem to ensure realistic diffusive behaviour (see Appendix A. 3. for technical details).

Coarse-grained simulations of entire bacterial genomes are computationally challenging, as typical genomes span ~ 5 Mbp, confined within micron-scale volumes. Any feasible simulation must then scale down the system. As our primary goal is to quantify the effects of geometric confinement, we identify the relevant dimensionless constant as *R*_*c*_*/R*_*g*_, *the ratio of confinement radius to the polymer’s unconfined gyration radius*. However, when considering kinetics, density of the system cannot be ignored, which is not an independent parameter. The polymer volume to confinement volume ratio, scales as: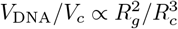.

To probe the role of geometric confinement, we simulated 2.4 kbp chains, confined within sphere of radius *R*_*c*_ = 0.3 − 1 times *R*_*g*_, where *R*_*g*_ ≈ 118.3 nm, which correspond to DNA volume fraction *V*_*DNA*_*/V*_*c*_ = 0.05 − 1.8%. Considering that simple bacteria show *V*_*DNA*_*/V*_*c*_ ≈ 1%, this choice of system size allows us to systematically vary the confinement radius, while avoiding biologically unrealistic high densities.

### B. Order Parameter for Tracking Sequence Pairing

To quantify the cumulative behaviour of pairing of homologous beads, we defined the following order parameter, at a given timestep *t*:

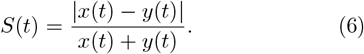

Here *x*(*t*) is the average distance between all ‘matching’ beads, and *y*(*t*) is the average distance between all nonmatching beads. For each timestep, we calculate the distance tensor *D*_*ij*_(*t*), the components of which indicate the centre-to-centre distance between bead *i* in one DNA molecule, and bead *j* in the other DNA molecule. We can write *x*(*t*) and *y*(*t*) explicitly in terms of *D*_*ij*_(*t*) as:

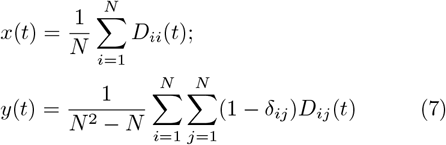

where *δ*_*ij*_ is the Kronecker delta. In the case of ideal, complete pairing of two overall homologous DNA, the distance between matching beads would be much less than the distance between non-matching beads. For such an extreme situation, *S* would have continuously evolved and eventually reached ~ 1. In contrast, in the case of two overall heterologous molecules, *S* would remain at ~ 0. Recent theoretical studies suggest, however, that pairing between long homologous sequences can never be fully complete due to thermal fluctuations, which give rise to persistent ‘bubbles’ [37], i.e. unpaired regions distributed along the contour length, in agreement with earlier experimental observations [9]). We expect to see this in simulation results.

### C. Counterion Condensation

To examine how counterion condensation affects the homologous pairing, we vary the parameter *θ* in our simulations, the fraction of phosphate charge neutralized by adsorbed cations. For a single DNA adsorbing cation, this parameter can be approximated with a Langmuir isotherm [11]:

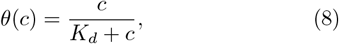

where *c* is the free cation concentration and *K*_*d*_ is the dissociation constant for binding to DNA. We consider two cases:

i. An idealized limit of full charge neutralization, *θ* = 1, corresponding to *c* ≫ *K*_*d*_.
ii. One that more realistically mimics conditions in bacteria, in which only a fraction of the DNA charge is compensated. To model this case, we use *θ* = 0.85. Assuming that magnesium (as the most abundant cation that condenses on DNA in prokaryotes) is the only specie that significantly adsorbes on DNA, using the Mg^2+^ dissociation constant *K*_*d*_ = 16.3 mM for approximately physiological monovalent ion concentration (100 mM Na^+^) reported by Ref. [11], we can relate *θ* = 0.85 to Mg^2+^ concentration of approximately 92 mM. This is consistent with the approximate total intracellular Mg^2+^ concentration in simple bacteria [39]. However, most Mg^2+^ is bound to phosphate-rich biomolecules, resulting in a free ion concentration on the order of mM. However, The dynamic nature of Mg^2+^ association, along with the presence of the polyamine spermidine, which binds strongly to DNA, makes *θ* = 0.85 a reasonable approximation.

## III. RESULTS

### A. Steady-state properties of paired homologous genes

Simulations were initialized by placing 25 central beads of each polymer at in 1-to-1 homologous register and randomising positions of the rest of the beads. This configuration was designed to mimic the state of the polymers after their first passage time, allowing us to focus specifically on the dynamics of pairing. Each simulation typically ran for approximately 0.5 ms until the system reached a steady state. At that point, each run resulted in one of two characteristic outcomes:

i. A 1-to-1 register of homologous sections (see Fig. 3a), in which each homologous section constantly “breathes” as it pairs and unpairs. This produces a set of paired and unpaired sections throughout the duplexes in each time snapshot.
ii. A kinetically stuck state in which a protruding duplex is caught in between two paired sections (see Fig. 3b). This latter type was more common for strong confinements, *R*_*c*_*/R*_*g*_ ≲ 0.4, for which the probability of homologous beads encountering each other and starting to pair at a different sight is increased.

**FIG. 3.**
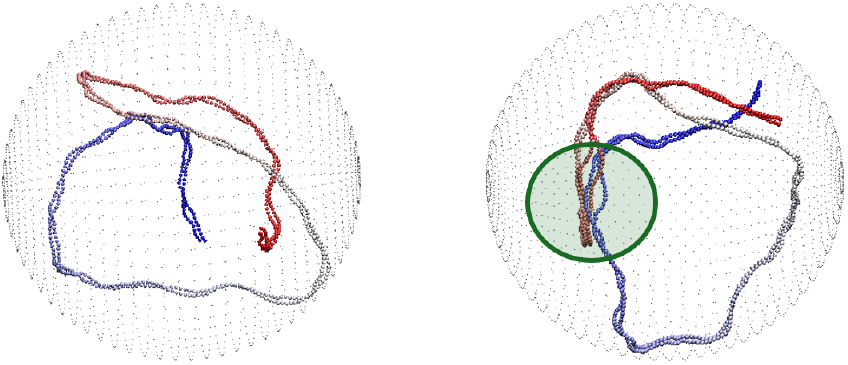
Pairing of two homologous DNA molecules: two types of steady state outcomes. Panels display the snapshots of simulation outcomes for two overall homologous DNA molecules. Each bead number (from 3’ to 5’ ends) is represented with a specific colour to distinguish different sections of homologous sequences from each other; each DNA duplex is painted by a colour-gradient from blue to red along its contour. Parameters used for both simulation runs were the same: *R*_*c*_*/R*_*g*_ = 0.7, *θ* = 0.85. The faint grey dots show the confining sphere. Outcomes: (a) Type 1: no kinetic traps, all beads are engaged in a periodic steady state ‘breathing’ from paired to unpaired local configurations and back. See movie for temporal evolution from initial state at https://youtu.be/xOvuRbSZvtQ. (b) Type 2: system is kinetically stuck due to protruding duplex going in between two paired regions. The highlighted green circle shows a problematic, stuck region where a duplex entered a looped section. See movie for temporal evolution from initial state at https://youtu.be/_D8-KT4FuJg.

Outcome (i) was reached directly or after first reaching outcome (ii) and then unpairing to reach a perfect to-1 register. This scenario was common for moderate confinement *R*_*c*_*/R*_*g*_ ≈ 0.5 −0.7, where a probability of encounter at a different sight is significant, but the duplexes have enough space to make these adjustments.

We quantified these steady states using the order parameter *S*(*t*), and report the time-averaged value over the *first* steady-state plateau, 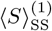, for fully homologous chains across confinement strengths and intrinsic charge compensation *θ*. When trajectories exhibited more than one plateau, i.e. first a kinetically stuck configuration and later the 1-to-1 register, we assigned ⟨ *S* ⟩_SS_ to the earlier (stuck) plateau (e.g. for *R*_*c*_*/R*_*g*_ = 0.7, *θ* = 0.85 in Fig. 4b). As confinement tightens, 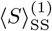 decreases, with a stronger reduction for *θ* = 0.85 than for *θ* = 1. This trend indicates a progressive shift from efficiently paired dynamics to kinetically trapped configurations, with trapping occurring more readily under weaker confinement when charge compensation is lower, and thus when the pairing strength is weaker.

**FIG. 4.**
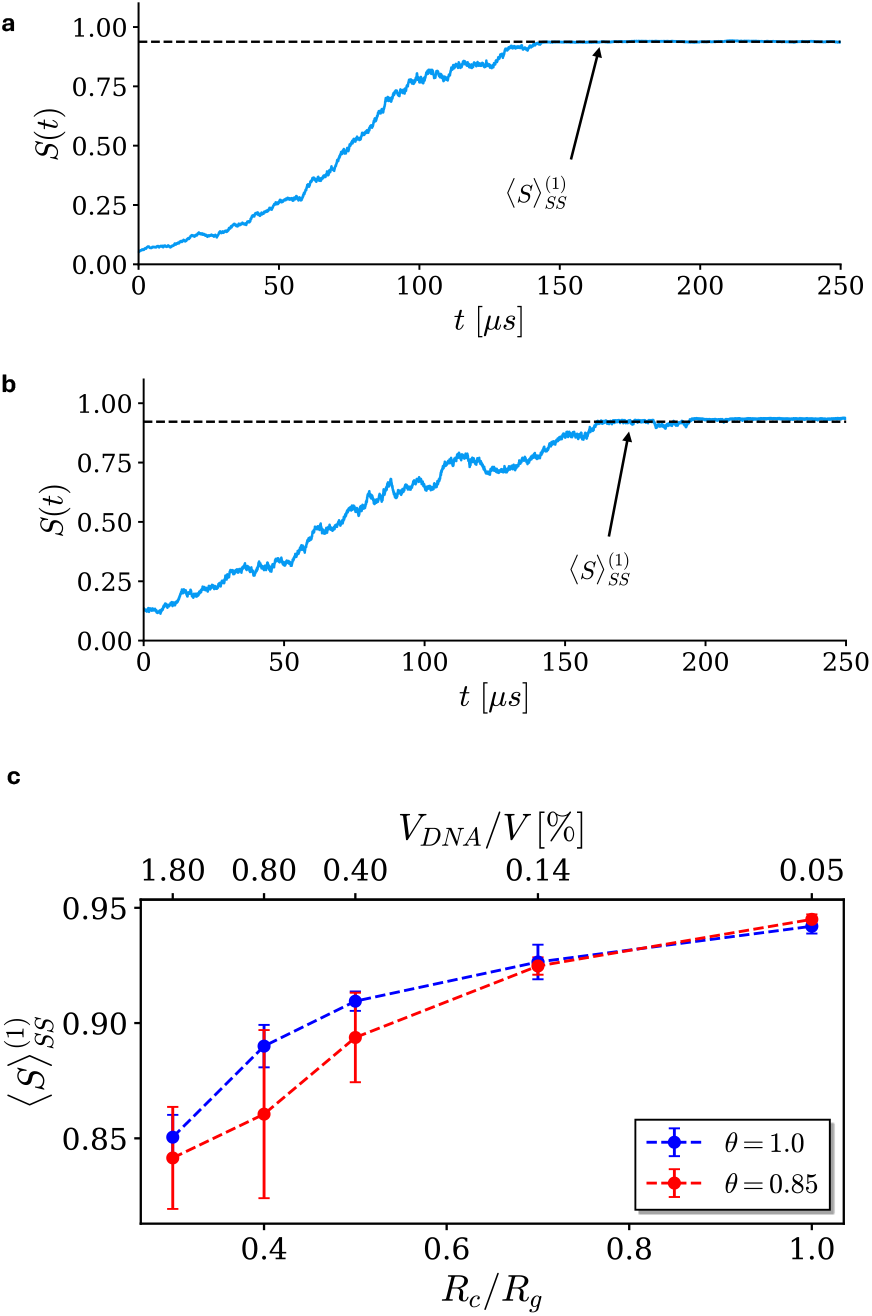
Time evolution of the order parameter at the first steady state 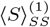; its dependence on confinement and fraction of charge compensated. (a) Time behaviour of the order parameter tracks pairing throughout the simulation. For the case of *R*_*c*_*/R*_*g*_ = 1.0 and *θ* = 1.0, order parameter converges to an equilibrium value, as a perfectly paired state is formed. (b) For the case of *R*_*c*_*/R*_*g*_ = 0.7 and *θ* = 0.85, order parameter reaches a first steady value, corresponding to kinetically stuck state, and then goes up to reach another steady state value, as a result of spontaneous unbinding and removing protrusion to obtain 1-to-1 register of homologous section. (c) The dependence of the steady state order parameter with confinement size for two chargecompensation levels, *θ* = 0.85 (red) and *θ* = 1.0 (blue).

For non-stuck system configurations, the distribution of distances in steady-state between homologous beads was recorded, see Fig. 5 (left) for the histograms. This distribution was used to analyse meaningful statistics of the system’s steady state properties, within a statistical physics framework. The height of each bin, approximates the probability *p*(*R*) of finding a two homologous beads at a distance *R*, and can be expressed as:

**FIG. 5.**
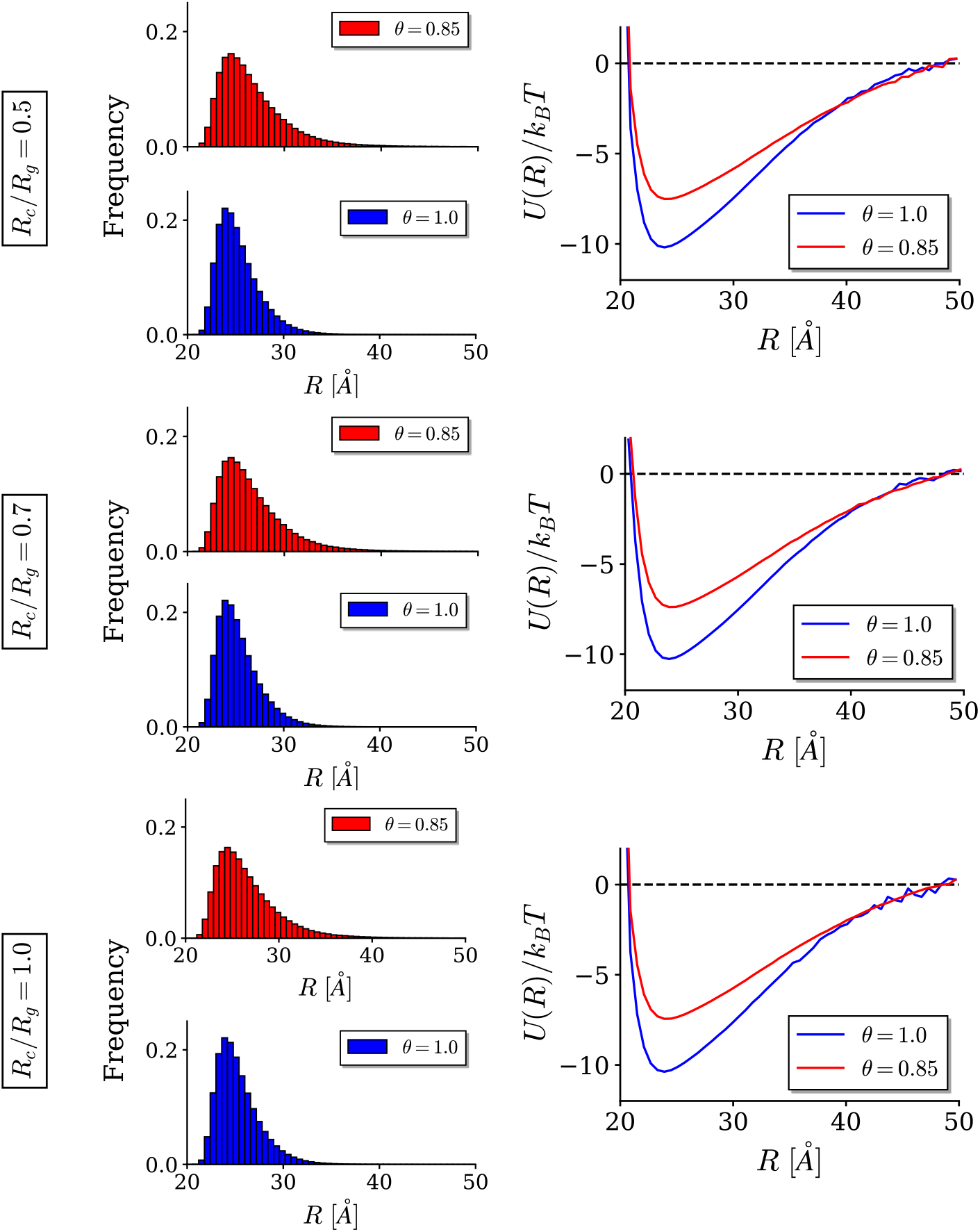
Distributions of distances between homologous beads and the resulting effective free-energy profiles. *Left column:* histograms representing distribution of distances between homologous beads recorded in steady state are shown for three confinement radii, *R*_*c*_*/R*_*g*_ = 0.5, 0.7 and 1, and indicated values of intrinsic charge compensation parameter *θ*. Statistics were gathered only for cases which achieved perfect pairing. *Right column:* Effective interaction potentials, *U* (*R*) for two charge-compensation levels, *θ* = 1.0 (blue) and *θ* = 0.85 (red), computed from their corresponding histograms by Eq. 10.

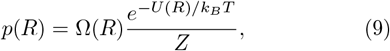

where *Z* is the partition function, Ω(*R*) is the multiplicity function (number of degenerate states) and *U* (*R*) is the effective energy between each two homologous beads which accounts for not only the electrostatic nonbonding interactions, but also the bonding (force-field) contributions between the beads alongside any effects from the confinement. The multiplicity function 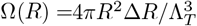, i.e. the ratio of the spherical shell volume associated with the specific bin position *R* and the cubed de-Broglie thermal wavelength, 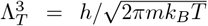, where Δ*R* is the width of the bin, *h* is the Planck constant and *m* is the mass of a bead.

The effective energy *U* (*R*) between two homologous beads is given by:

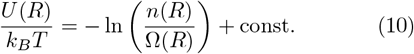

To effectively remove the constant from Eq. 10 we simply set *U* (*R* → ∞) → 0, i.e. we vertically shift the entire curve such that the tail ends at zero. Results for *U* (*R*) for different (non-stuck) cases are shown in Fig. 5 (right). Comparing the *U* (*R*) in Fig. 5 with the result in Fig. 2, reveals how collective behaviour of the polymer beads impacts the interactions between two homologous beads. The well is wider and shallower for *θ* = 0.85, reflecting how smaller charge compensation results in less tightly bound states.

### B. Kinetics of homologous pairing: evolution to steady state

The characteristic time to pairing, *t* = *τ*, was taken to be the time at which the order parameter converged to its first steady state value, 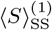 (see Fig. 4a and Fig. 4b). The pairing time as a function of confinement for two different intrinsic charge compensations *θ*, is shown in Fig. 6a. As *R*_*c*_*/R*_*g*_ decreases from the weakconfinement limit (*R*_*c*_*/R*_*g*_ = 1), the pairing time drops sharply, demonstrating that geometric confinement accelerates pairing by restricting configurational exploration. However, once confinement becomes sufficiently tight, the pairing time reaches a plateau. Combined with data in Fig. 4c, this behaviour reflects the onset of protrusion formation, which restricts molecular motion and counteracts further acceleration. This plateau occurs at larger *R*_*c*_*/R*_*g*_ for *θ* = 0.85. In this regime, pairing is slower, so homologous segments distant from the initially paired region have more time to search and interact, increasing the likelihood of protrusion formation and thus shifting the plateau to weaker confinement.

**FIG. 6.**
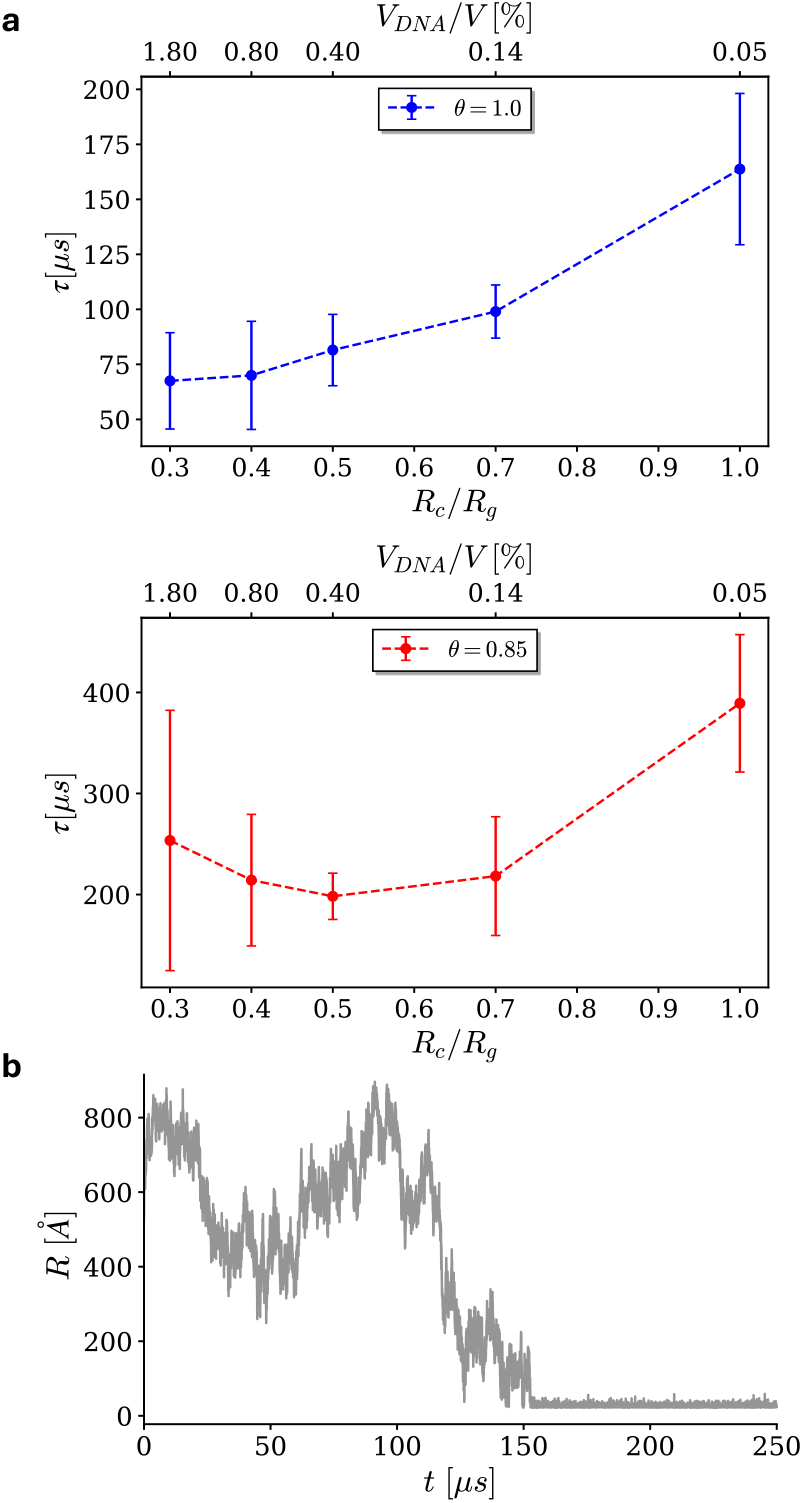
Pairing kinetics. (a) The homologous pairing time, *τ*, as a function of confinement, for *θ* = 0.85 (red) and *θ* = 1.0 (blue). Points were obtained as a mean over four independent repeats and their error bars display standard deviation for the different repeats. (b) Time evolution of centre-to-centre separation between randomly selected couple of homologous beads, plotted for *θ* = 0.85 and confinement *R*_*c*_*/R*_*g*_ = 0.7.

The temporal evolution of the distance between two specific homologous beads is shown in Fig. 6b. Once in the paired state, we see that the beads exhibit some steady-state “breathing mode” in which the distance between them fluctuates over time. Thus, while the order parameter gives a rough estimate of the pairing status, it lacks the ability to predict the amount of paired homologous beads. Therefore, we define two homologous beads to be paired when they are at a surface-to-surface separation equal or lower than Debye length, *λ*_*D*_ = 7 Å as a length scale for electrostatic interactions. Given this definition, we calculated the time evolution of the fraction of homologous paired beads, *n*(*t*) as each simulation progresses, shown in Fig. 7b (grey points).

**FIG. 7.**
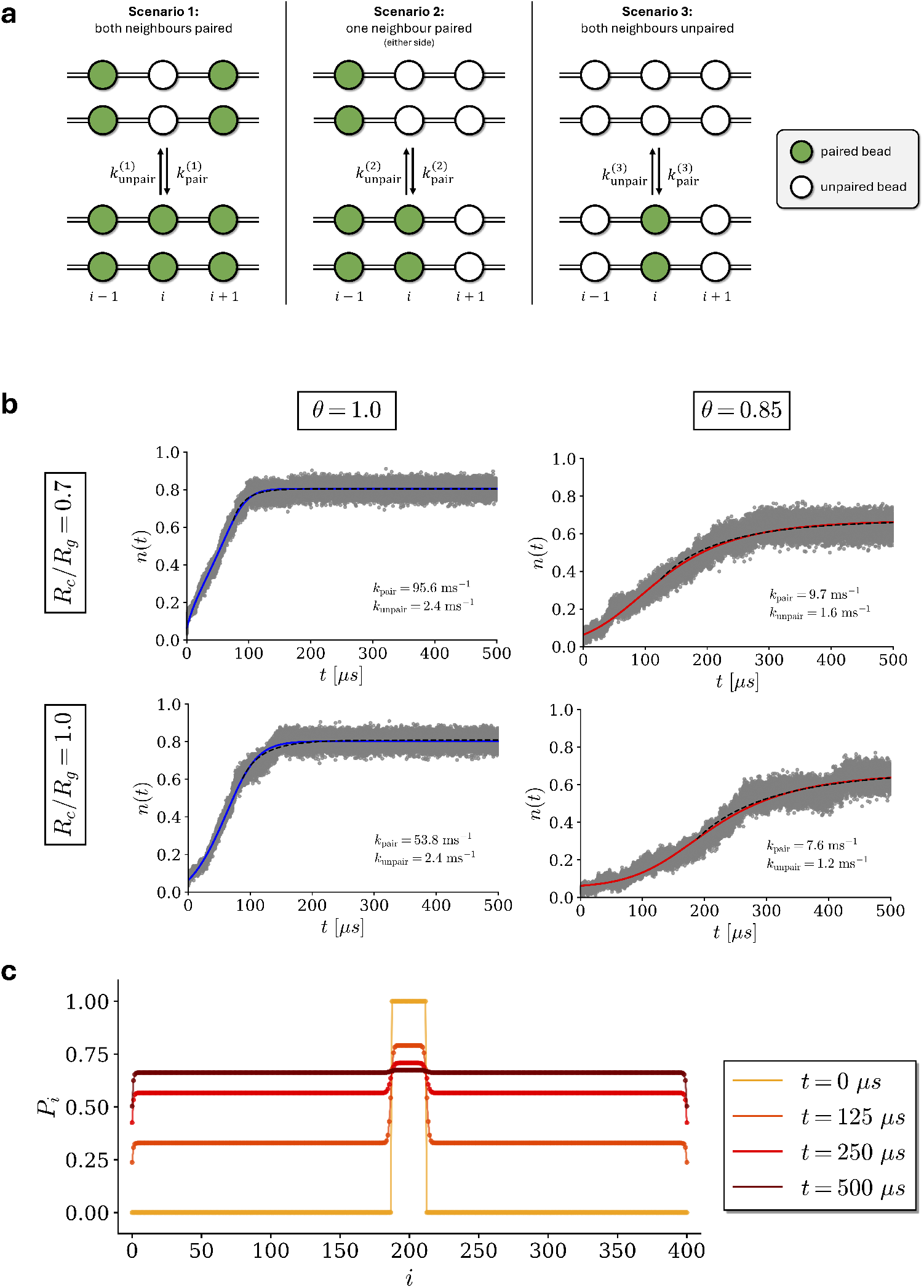
The kinetics of homologous gene pairing. (a) Schematic of the model for homologous gene pairing used in *Detailed Master Equation*, which takes into account correlations with the nearest neighbours. (b) The progression of pairing is shown for various cases of confinement and charge compensation. Raw simulation data are shown in gray. The fits to the *Simple Kinetic Model*, expressed by Eq. 11, are shown as dashed lines, with the fitting parameters, *k*_pair_ and *k*_unpair_, shown in each panel. The fits to the *Detailed Master Equation*, expressed by Eq. 12, are show as blue (for *θ* = 1.0) and red (for *θ* = 0.85) solid lines. The fitting parameters are displayed in Table I. (c) *P*_*i*_ as a function of bead index, *i*. Example plot for *θ* = 0.85, *R*_*c*_*/R*_*g*_ = 0.7 at four representative times.

#### Simple Kinetic Model

To characterise the pairing and unpairing dynamics of beads, we derived a simplistic kinetic theory in which we consider two rates; *k*_pair_, the rate of formation of a pair of beads and *k*_unpair_, the rate of dissociation of a pair of beads. Constructing the corresponding rate equation for *n*(*t* − *t*_0_), fraction of beads paired at times larger than empirically chosen time *t*_0_, and solving it analytically (see Appendix B.1 for details), we obtain:

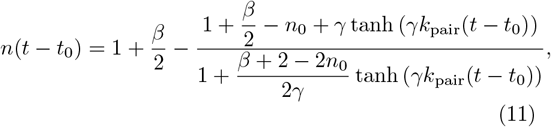

where *β* = *k*_unpair_*/k*_pair_ and 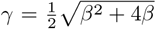. This simple model is expected to provide a reasonable description of the system near equilibrium, but less-accurate description in the stages where the system is far from equilibrium. Fitting Eq. 11 to the simulation data (dashed lines in Fig. 7b),this simple model gives qualitatively reasonable, but not ideal fits (dashed curves in Fig. 7b), as it fails to reflect correlations between the beads within each chain.

#### Detailed Master Equation

The simple kinetic model above only considered the evolution of the global paired fraction *n*(*t*), and thus neglects correlations along the chain. In our simulations, the pairing/unpairing of beads is locally cooperative: beads within a paired domain are stabilised by their neighbours, whereas beads at domain boundaries and at the ends of the chain are more susceptible to unpairing, leading for example to fraying at the end of the paired dsDNA chains. Collapsing these distinct behaviours into a single pair of effective rates as in the simple kinetic model above necessarily averages over local environments and therefore cannot reproduce the detailed shape of *n*(*t*) at early stages of pairing.

To capture this, we introduce a nearest-neighbour master equation for the time-dependent probability that a bead *i* along the chain is paired, *P*_*i*_(*t*). The rate of change of *P*_*i*_ depends on whether its neighbours (*i* − 1 and *i* + 1) are paired, leading to three local environments: (1) both neighbours paired, (2) exactly one neighbour paired, and (3) both neighbours unpaired (see Fig. 7a). Each environment *s* ∈ {1, 2, 3} has its own pair-formation and dissociation rates, 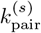 and 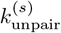, giving:

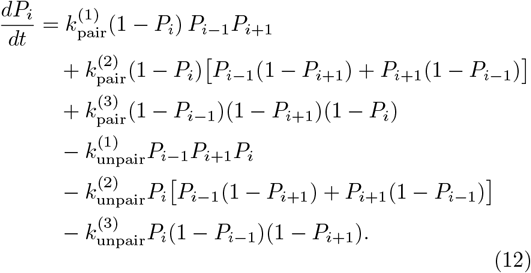

The first three terms represent pair formation in the three neighbour configurations, while the last three represent dissociation in their corresponding configurations. We solve this system numerically and compare to simulations via the mean paired fraction *n*(*t*) ≈ (1/*N*) ∑ _*i*_ *P*_*i*_(*t*). To reduce the number of independent fitting parameters, we impose an approximate detailed-balance constraint within each local environment, relating 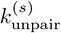 to 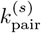 via equilibrium pairing probabilities (see Appendix B.2). The resulting fits (solid curves in Fig. 7b) capture fine structure in *n*(*t*) that is missed by the simple model, consistent with locally cooperative pairing/unpairing along the chain. Interestingly, as the system approaches equilibrium, the pairing probabilities for individual beads, *P*_*i*_ (obtained by numerically solving Eq. 12 with parameters from Table I), converge to a uniform value (see Fig. 7c for *θ* = 0.85, *R*_*c*_*/R*_*g*_ = 0.7). The exception is the terminal beads, which exhibit lower pairing probabilities, reflecting their increased tendency to fray. However, since these beads account for only a small fraction of the chain, the *Simple Kinetic Model* is expected to provide a reasonable description of the system near equilibrium.

**TABLE 1.** Rate constants for pairing and unpairing in different modes. The rate constants are presented for various cases of confinement and charge compensation and were obtained as fitting parameters from fitting Eq. 12 to curves in Fig. 7b.

| | $\theta = 1.0$ | | $\theta = 0.85$ | |
| --- | --- | --- | --- | --- |
| | $R_c/R_g = 0.7$ | $R_c/R_g = 1.0$ | $R_c/R_g = 0.7$ | $R_c/R_g = 1.0$ |
| $k_{\text{pair}}^{(1)} [\text{ms}^{-1}]$ | 124.3 | 76.8 | 13.6 | 7.5 |
| $k_{\text{unpair}}^{(1)} [\text{ms}^{-1}]$ | 13.8 | 8.5 | 2.1 | 1.2 |
| $k_{\text{pair}}^{(2)} [\text{ms}^{-1}]$ | 26.7 | 23.4 | 13.3 | 10.8 |
| $k_{\text{unpair}}^{(2)} [\text{ms}^{-1}]$ | 12.6 | 11.0 | 9.6 | 7.8 |
| $k_{\text{pair}}^{(3)} [\text{ms}^{-1}]$ | 15.3 | 4.1 | 1.5 | 0.3 |
| $k_{\text{unpair}}^{(3)} [\text{ms}^{-1}]$ | 36.8 | 9.9 | 7.2 | 1.4 |

## IV. DISCUSSION

Our study provides evidence that confinement - a fundamental physical feature of cellular environments directly influences the kinetics and outcomes of homologous DNA pairing, highlighting the importance of spatial organisation in genome dynamics [40–42]. Cells naturally exploit compartmentalisation to create distinct microenvironments that enhance biochemical efficiency, as seen in the DNA-rich nucleoid and nucleoli compartments and prokaryotes and eukaryotes [42, 43]. These structures are thought to locally modulate reaction rates and coordinate genomic processes; they may also potentially facilitate homologous recombination. Our findings suggest that global geometric confinement, even in the absence of proteins, can promote homologous pairing, suggesting that cells may inherently take advantage of their physical architecture to regulate genome activity.

On a molecular level, our simulation results reinforce that DNA’s capacity for homology recognition is an intrinsic physical feature, modulated by sequence-dependent structural attributes and electrostatic interactions that become biased under confinement. The emergence of an optimal pairing regime, hints that evolution may have selected for cellular sizes that naturally expedite essential genomic processes [44, 45]. Such observations parallel experimental findings where restricted volumes in biomolecular condensates enhance reaction kinetics by reducing the configurational search space [46, 47]. The dependence of optimal confinement on the fraction of phosphate charge neutralized suggests that cell size optimisation is shaped by a complex interplay of intracellular conditions, for example the concentration and identity of DNA-condensing cations. These physicochemical effects provide an additional argument for the wide diversity of bacterial cell sizes observed in nature [48]. Our findings may also point to potential fundamental size limit in living systems [49] where excessive confinement impairs DNA pairing efficiency, suggesting that cells and organelles cannot be too small without compromising essential genomic dynamics.

The analysis of homologous pairing kinetics provides important insights into the pairing pathways of long chains. The two kinetic models presented in the main text account for pairing in regions near-equilibrium and far from equilibrium. The more sophisticated kinetic approach based on a detailed Master Equation, accounts for transient configurations and can therefore be used – within some reservations – to describe the behaviour over the entire time domain. While this fit is not exact, it is quite accurate despite the substantial scatter of the data caused by fluctuations. Importantly, the more refined model ultimately validates the two-state description of the system in the simpler kinetic Model; it predicts that the paired probability as a function of bead index approaches a constant value - hence, the nonlinear contributions cancel out, leaving only the terms captured by the two-state model.

These insights hold significant promise for synthetic biology and artificial cell technologies [50, 51]. In bottomup approaches to synthetic cell construction, the incorporation of geometric confinement—through vesicles, coacervates, or droplet compartments—can be strategically harnessed to direct homologous DNA interactions and recombination events [52]. By embedding physical constraints into cell-free systems, it may be possible to recapitulate complex genome-scale functions without relying on the full complement of cellular protein machinery.

## V. CONCLUSIONS

This work establishes that homologous gene pairing is accelerated by geometric confinement imposed by a spherical boundary. The timescale of homology search is critically governed by the dimensions of the confining compartment: excessive spatial freedom prolongs the search, while excessive restriction impedes recognition kinetics.

These findings reveal the existence of an optimal confinement regime that maximizes the efficiency of homology search. This confinement, varies with fraction of charge compensated. It is plausible that biological systems leverage this physical principle to reconcile rapid genetic recombination with other essential cellular constraints.

## VI. ACKNOWLEDGEMENTS

The authors are thankful to Marco di Antonio, Geoff Baldwin, Fernando Bresme, Ming Chen, Claudia Danilowicz, Ralph Holden, Lorenzo di Michele, Mara Prentiss and John Seddon for useful discussions throughout this work. This study was part of a project supported by the Leverhulme Trust, Grant #RPG-2022-142. JGH is supported by a Novo Nordisk Postdoctoral Fellowship in partnership with the University of Oxford.

## VII. AUTHOR CONTRIBUTIONS

AAK and EH conceived the project. EH, JGH, AR, GO, and AAK developed methodology, NES, JGH and YX performed simulations and curated the data, EH, JGH, NES, YX, AS, GO, and AAK analysed the data, NES, JGH, and YX performed their visualization. EH, AAK led the writing of the initial draft, with drafting and revision contributions from JGH; NES developed its submitted version, with input from all authors. AS provided key conceptual suggestions during revision, informed by ongoing experimental work to test the paper’s predictions. All authors participated in review and editing of the paper and approved the final manuscript. AAK supervised the project and, with YE, acquired funding.

## Appendix A

**Simulation Methods**

### 1. Simulation Setup

The coarse-grained molecular dynamics simulations of two DNA molecules confined within a spherical cavity, were performed employing the Largescale Atomic/Molecular Massively Parallel Simulator (LAMMPS) software package [53]. DNA molecules were modelled using bead-spring polymers, with each bead representing approximately 20 Å section (approximately six base pairs) of double-stranded DNA.

Initial polymer conformations were systematically generated using custom Python scripts. 25 beads in the middle of each polymer were place in a straight line at the centre of the confining sphere at a separation of 23.9 Å, corresponding to the minimum of the potential well (see Fig. 5). The rest of the beads were placed sequentially in the confining sphere using a self-avoiding random walk, with collinear segments of 50 beads (matching the Kuhn length). Trial positions that exceeded the confinement radius or overlapped with any other bead were rejected and new random directions were sampled.

Confinement was imposed by a repulsive radial force acting near the boundary of the spherical volume, derived under the Derjaguin approximation to reflect the steric and electrostatic repulsion experienced by DNA near confining surfaces. This potential ensured that polymer beads remained within the spherical domain while allowing physically realistic proximity to the boundary (see main text for more details). Both bonded (polymer) and non-bonded (electrostatic) interactions were approximated through tailored potentials reflecting the physical electrostatic characteristics of DNA in solutions of the physiological electrolyte concentration, as described below.

Temperature was maintained at 37^°^C using Langevin dynamics thermostatting combined with NVE integration, employing a damping coefficient consistent with the DNA hydration environment. See below for estimation of this parameter. Equations of motion were integrated using a timestep of 500 fs.

Each simulation consisted of 500 million timesteps, corresponding to a total simulation time of 250 *µs*. If the order parameter (defined in the main text, Eq. 6) had not indicated convergence in pairing by the end of the simulation, the run was restarted using the final configuration. Computational runs were executed using parallel processing via Message Passing Interface (MPI), distributed across 64 CPU cores. Typical simulation runtimes for each 500 million timestep run were approximately 30 hours, enabling extensive parameter-space exploration within computationally tractable durations.

#### 2. Interactions between beads in the same DNA chain

To maintain the mechanical properties of a DNA chain, we implemented a set of interactions between neighbouring beads of the same polymer. For resistance to length deformations, a 12-6 Lennard-Jones (LJ) interaction was implemented between each two consecutive beads,

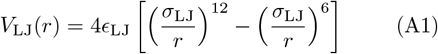

where *r* is the centre-to-centre distance between the two consecutive beads and {*ϵ*_LJ_, *σ*_LJ}_ is the set of LJ parameters estimated to reproduce the mechanical properties of DNA. Taking into consideration the radius of DNA, we set the minimum distance at *r*_min_ = 2.1 nm, from which we determined *σ*_LJ_ = 2^−1*/*6^*r*_min_ ≈ 2.35 nm. To set *ϵ*_LJ_, we expanded Eq. A1 to second order:

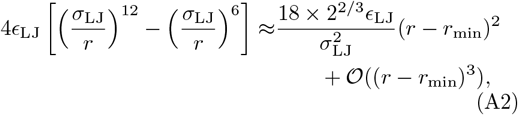

and compared it with the elastic energy stored in a stretched rod characterised with the same Young modulus as that of a DNA molecule, *E* ≈ 1.1 × 10^−1^ nN nm^−2^, to obtain *ϵ*_LJ_ ≈ 233.5 *k*_*B*_*T*, where the temperature is set to 37^°^C.

To take into account resistance against bending, a cosine potential of the form:

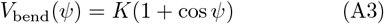

was assigned for each two consecutive sections of beads, i.e. for each triplet of beads {*i* − 1, *i, i* + 1}. Here, *ψ* is the angle between centre-to-centre line of beads {*I* − 1, *i* }, to that of beads {*i, i* + 1}. When the polymer is locally straight, we have *ψ* = *π*, which gives zero bending penalty. The *K* coefficient was set by expanding Eq. A3 for *ψ* around the minimum energy to second order:

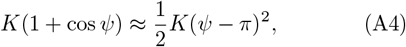

**FIG. A1.**
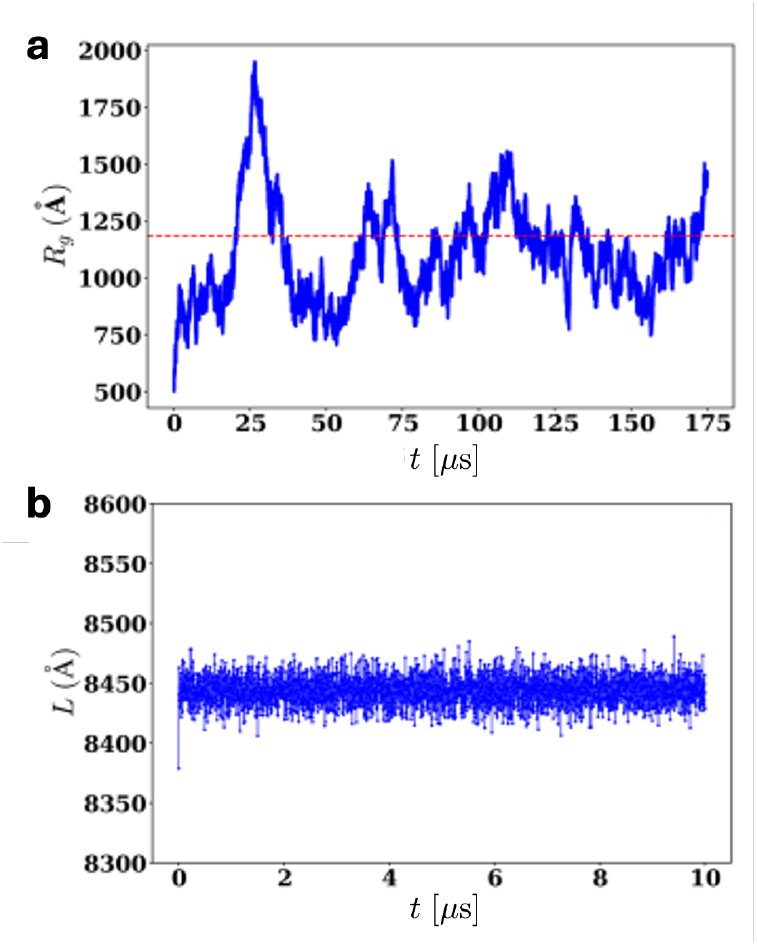
A simulation of a single 2.4 kbp dsDNA chain in free space. (a) the time evolution of the gyration radius. Dashed line shows the theoretical prediction (1183.2 Å). (b) The time evolution of the total arc length of the chain. The value of the total arc length is shown to be practically a constant, albeit about 3% larger than the expected value due to the interspace separation of 1 Å between the surfaces of each two consecutive beads, and thermal expansion of the chain.

and comparing it with the elastic energy stored in a torsional spring having the same flexural rigidity as that of a DNA molecule, which came out as:

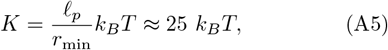

where ℓ_*p*_ ≈ 50 nm is the persistence length of DNA in normal physiological conditions.

To check if these interactions were implemented correctly, we ran the simulation for a single 2.4 kbp long chain in free space, which is essentially a periodic simulation box with dimensions much larger than the theoretical gyration radius of the polymer. We recorded the total arc length, *L*, and gyration radius, *R*_*g*_, as a function of time. The gyration radius was calculated according to its definition,

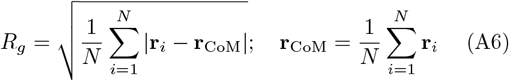

where **r**_*i*_ is the position of the centre of the *i*-th bead, and **r**_CoM_ is the centre of mass of the polymer.

The results for *R*_*g*_ and *L* are shown in Fig. A1, which indicate that the arc length of the chain is close to invariant under thermal fluctuations, as appropriate, and the mean steady state gyration radius fluctuates about the value predicted by standard polymer theory, i.e.,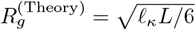 is the Kuhn length of the polymer.

#### 3. Homology Recognition Interaction Coefficients

The interaction between homologous beads at centreto-centre distance *R* is described by Eq. 4 in the main text, inspired by the theory of electrostatic interaction of helical molecules [32]. Here, the interaction is written in the form of an expansion in helical harmonics, *a*_*n*_(*R*), where we consider the first few dominant terms. For a local response of the electrolyte solution in which the DNA are solvated (*ε* = 80), these are given by:

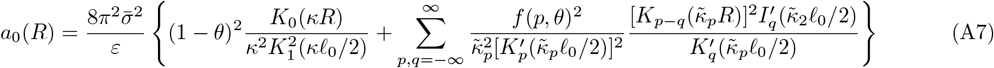

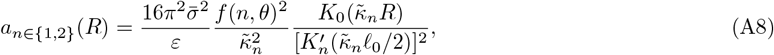

where 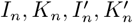 are the modified Bessel functions and their derivatives, *κ* is the inverse Debye length evaluated at physiological conditions, 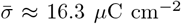 is the average surface charge density of the uncompensated phosphate charge, 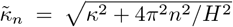 is the *n*-th harmonic effective inverse screening length, where *H* is the helical pitch of DNA. Lastly,

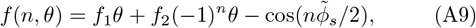

where *f*_1_ and *f*_2_ = 1 − *f*_1_ are the fractions of counterions adsorbed to the minor and major grooves respectively, and 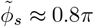 is the azimuthal width of the minor groove.

We assumed a type of salt that dominantly adsorbs to the major groove, such as Ca^2+^ or Mg^2+^, for which we set *f*_1_ = 0.2 and *f*_2_ = 0.8 in all simulations.

Assuming that the rotation of the beads is instantaneous compared to diffusive properties of the polymer, the ‘relative azimuthal orientation’ *δϕ* between two beads can be taken as the path that minimises the electrostatic interaction energy:

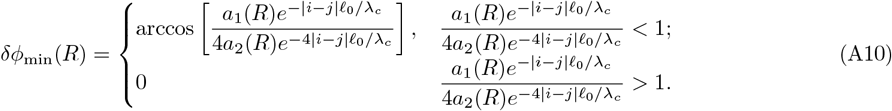

**FIG. A2.**
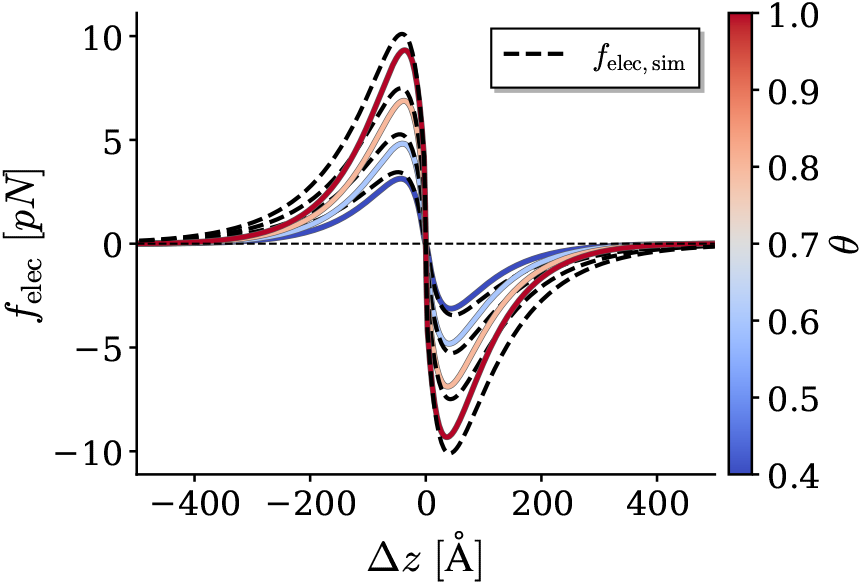
Exact Theory vs. Simulation Approximation The electric force, *f*_elec_ = − ∂U_elec_*/*∂Δ*z*, as a function of the axial shift, Δ*z*, stemming from: 1. the full electrostatic potential, Eq. 2, (solid curves), and, 2. the approximated version, Eq. 4, used in the simulation (dashed), where Δ*z* = *i j* ℓ_0_ was taken to be continuous for visualisation. Curves were plotted for a range of different values of intrinsic charge compensation, *θ*.

The exact expression for the electrostatic energy of two beads originating from different DNA molecules depends on the interaction length, *L*, in a non-linear fashion as shown in Eq. 3. It is computationally expensive to quantify the interaction length for interacting sections and produce, accordingly, a time-changing electrostatic profile. Therefore, we prescribed an approximated long interaction length (*L* ≫ *λ*_*c*_) forcefield for the beads. The electric force *f*_elec_ = − ∂U_elec_*/*∂Δ*z* that such a potential creates is a very good approximation for small shifts Δ*z* ≪ λ_c_, which is self-consistent with the final outcome of the simulation in which the genes that pair-up do so with a very small shift Δ*z* ≪ λ_c_. Furthermore, this approximation turns out to be quite accurate even for large Δ*z* shifts, as depicted in Fig. A2.

#### 4. Langevin Dynamics

At each timestep, the position of each bead, **r**, was displaced according to the Langevin equation, which reads as:

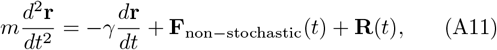

where *m* is the mass of the bead, *γ* is the hydrodynamic drag coefficient of the bead in creeping flow, **F**_non−stochastic_ is the sum of mechanical, electrostatic and hydration forces acting on the bead, and **R** is a random force taken from a Gaussian distribution which satisfies the following autocorrelation:

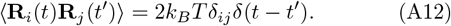

The hydrodynamic drag coefficient was approximated using Stokes’ law:

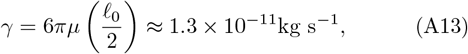

where *µ* ≈ 0.7 × 10^−3^ Pa s is the dynamic viscosity of water in 37^°^C and ℓ_0_ = 2 nm is the diameter of a single bead in the DNA chain. The molar mass of each bead is approximated to be 3960 g mol^−1^, leading to a Langevin damping time of *τ* = *m*_bead_*/γ* ≈ 505 fs. By estimating this parameter for water our simulation replicates real time evolution.

## Appendix B

**Kinetic Models**

### 1. Simple Kinetic Model

Here we describe the kinetics of homologous pairing, which treats homologous beads independent of their neighbours, in the case when the system does not get ‘trapped’ on the way to reaching equilibrium. After initiating a segment of 25 beads in parallel juxtaposition, the time evolution of the fraction of paired homologous beads, *n*(*t*) = *N*_paired_(*t*)*/N*, which is complementary to the fraction of *unpaired* homologous beads, can be approximated as :

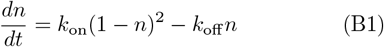

where *k*_on_ and *k*_off_ are the kinetic coefficients for, respectively, creating or destroying the pairing between two homologous beads. Applying initial conditions, *n*(*t*_0_) = *n*_0_, where *t*_0_ is empirically chosen time, when the kinetics start to follow the *Simple Kinetic Model*, Eq. B1 is solved analytically by separation of variables:

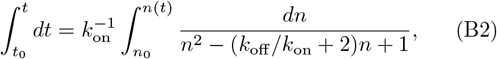

which gives

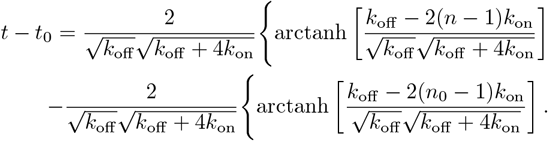

Rearranging the right-hand side of this equation to solve for *n*(*t*), and defining *β* = *k*_off_ */k*_on_, we obtain Eq. 11 of the main text. At small times, 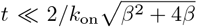, Eq. 11 gives a linear rise:

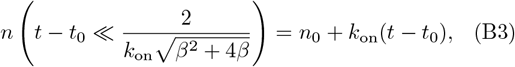

and at long enough times 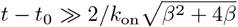, the number of paired sites approaches its steady state value:

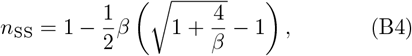

which gives *n*_SS_ → 0 for *β* → ∞, and *n*_SS_ → 1 for *β* → 0, as required.

#### 2. Detailed Master Equation

##### a. Model Formulation

To incorporate correlations along the chain, we track the probability that each bead is paired, *P*_*i*_(*t*). The key modelling assumption is nearest-neighbour dependence,the kinetics at bead *i* depends on whether beads *i* − 1 and *i* + 1 are paired. This introduces three local environments for bead *i*:

1. both neighbours are paired,
2. exactly one neighbour is paired,
3. both neighbours are unpaired.

We assign each environment *s* ∈ {1, 2, 3} its own pairformation and dissociation rates 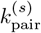 and 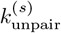. By expressing the probability of each neighbour configuration in terms of *P*_*i*−1_, *P*_*i*_ and *P*_*i*+1_, the overall evolution of *P*_*i*_(*t*) can be written as gains from pair formation minus the losses from dissociation, yielding Eq. 12 in the main text. For compactness we let *P*_*i*_ ≡ P_i_(*t*).

##### b. Detailed-Balance Approximation

Directly fitting all six rates (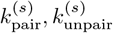 for *s* =1,2,3) can be poorly constrained. We therefore reduce the number of independent parameters by imposing an approximate detailed-balance condition within each local environment *s*. At equilibrium, conditional on being in scenario *s*, we can therefore write::

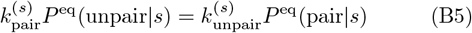

where *P*^eq^(unpair | *s*) and *P*^eq^(pair|*s*) are equilibrium probabilities that two homologous beads are unpaired and paired providing that scenario *s* occurs. Applying Eq. B5 to Eq. 12 eliminates 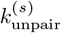 for each *s*, thereby reducing the number of free kinetic parameters used in the fitting, resulting in:

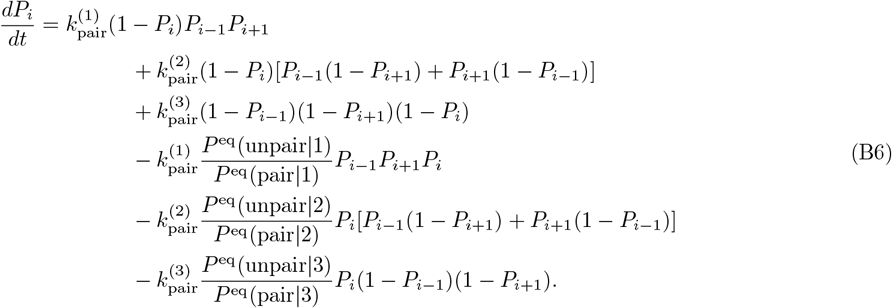

##### c. Numerical solution and boundary conditions

We solve Eq. B6 for *P*_*i*_(*t*) numerically using a fourth-order Runge-Kutta (RK4) integrator. Initial conditions match our protocol for MD simulations: *P*_*i*_(*t* = 0) = 0 for *i* ∈ [1, 187] and *i* ∈ [213 − 400], and *P*_*i*_(*t* = 0) = 1 for *i* ∈ [188, 212]. To implement open boundaries and allow end effects such as fraying, we introduce two ‘ghost’ beads at *i* = 0 and *i* = *N* + 1 with fixed pairing probability *P*_0_(*t*) = *P*_*N*+1_(*t*) = 0. This prevents the “bothneighbours-paired” environment from occurring at terminal beads and naturally incorporates reduced stability at the chain ends.

##### d. Connection to observables and fitting

To compare the dynamics modelled in this master equation to simulation, we approximate the instantaneous paired fraction by the mean of bead pairing probabilities:

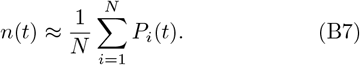

For a given parameter set, we integrate the master equation to obtain *n*(*t*) and fit to the simulation times series data. The resulting curves are shown as solid lines in Fig. 7, and fitted parameters in Table I.

